# Multimodal human and pet cues intensify wildlife fear responses

**DOI:** 10.64898/2026.05.13.725053

**Authors:** Kota Hirobe, Masayuki Senzaki

**Affiliations:** Graduate School of Environmental Science, Hokkaido University, Sapporo, Hokkaido, Japan; Faculty of Environmental Earth Science, Hokkaido University, Sapporo, Hokkaido, Japan

**Keywords:** alert distance, *Cervus nippon yezoensis*, domestic dog, fear of humans, flight initiation distance, human-wildlife coexistence, multimodal cue, sensory cue

## Abstract

Fear of humans can drive persistent changes in wildlife behavioural and life-history traits, with cascading effects on entire ecosystems. Human multimodal cues and pet cues may influence such fear, yet no study has tested how wildlife fear responses change when human acoustic cues and pet visual and acoustic cues are added to human visual cues. Filling this gap is important for managing outdoor activities for humans and pets while conserving wildlife. Here, using dogs as a representative pet, we tested the effects of human and dog visual and acoustic cues on fear responses of sika deer (*Cervus nippon yesoensis*) across an ∼800 km^2^ area in northern Japan, using alert distance (AD) and flight initiation distance (FID). We measured both distances with an approaching surveyor alone and with additional cues. Analyses with 266 observations showed that AD was estimated at 80.0 m with the human visual cue alone, and dog barking increased AD by 18.4 m. FID was estimated at 57.1 m with the human visual cue alone, and the human voice and the dog decoy increased FID by 11.3 m and 8.5 m, respectively. These results demonstrate that both human multimodal cues and pet cues can increase wildlife fear responses, suggesting that dog walking may expose wildlife to concurrent predator-associated cues more consistently than occurs under natural conditions. Our finding that adding human acoustic cues increased FID, contrary to previous studies, suggests that animals may adjust the relative weighting of sensory cues depending on predator species. These findings highlight the importance of considering the combined effects of human and pet cues when managing recreation in sensitive areas, favouring targeted measures over blanket restrictions.

---

Humans have caused wildlife declines and extinctions through land-use change, hunting, advances in killing technology, domestication, and other factors (Andermann et al., 2020; Barnosky et al., 2004; Cosentino & Gibbs, 2022; Darimont et al., 2015; Leveau, 2021; Uchida et al., 2019). Among these impacts, fear of humans is a key driver of persistent changes in foraging, reproduction, space use, and diel activity patterns (Gaynor et al., 2018; Lasky & Bombaci, 2023), with cascading effects on the structure and functioning of entire ecosystems (Smith et al., 2024; Suraci et al., 2019). Understanding how wildlife fear responses to humans manifest is, therefore, essential for developing context-specific management strategies that mitigate these impacts (Gaynor et al., 2021).

In ecological literature, fear of predators, including humans, can be defined as an animal’s conscious or unconscious perception of predation risk (Brown et al., 1999; Gaynor et al., 2021). Specifically, wildlife perceives predator cues (e.g., visual, acoustic, and olfactory cues), evaluates the risk of injury or death (Gaynor et al., 2021; Jones et al., 2024), and responds via anti-predator behaviours such as escape, often accompanied by physiological stress (Gaynor et al., 2021). The types and combinations of these predator cues can influence response magnitude, and simultaneous multimodal cues can amplify perceived predation risk, eliciting stronger anti-predator responses (Jones et al., 2024; Mukherjee et al., 2024; Munoz & Blumstein, 2012). For example, Jones et al. (2024) reported in a meta-analysis that prey show significantly larger behavioural changes to multimodal than to single predator cues. However, no study has explicitly compared wildlife fear responses to humans under multimodal versus unimodal cues, despite humans’ role as “super- predators” capable of intimidating even apex predators (Darimont et al., 2015; McGann et al., 2024).

Additionally, human impacts on wildlife, such as fear responses, may vary with human outdoor activity patterns (e.g., group size and whether people bring pets) (Banks & Bryant, 2007; Gómez-Serrano, 2021; Miller et al., 2001). Walking with large pets, such as dogs (*Canis lupus familiaris* Linnaeus) in particular, may heighten fear responses to humans, either because dogs chase or attack wildlife (Ritchie et al., 2013; Taylor-Brown et al., 2019) or are perceived as threatening (Banks & Bryant, 2007; Gómez-Serrano, 2021; Greenwell & Dunlop, 2023). This amplification may vary with the types and combinations of cues animals perceive (Jones et al., 2024; Munoz & Blumstein, 2012), including visual cues of humans and pets and acoustic cues such as human conversation and pet barking (Jones et al., 2024; Munoz & Blumstein, 2012). Although outdoor activities involving pets can negatively affect wildlife, many people continue to visit natural areas with pets (Packer et al., 2024; Westgarth et al., 2021), partly because nature-based walking benefits both human and pet health and well-being (Packer et al., 2024). Understanding how fear responses to humans with pets change with the types and combinations of perceived cues is, therefore, crucial for managing human outdoor activities while conserving wildlife.

Here, we examined how wildlife fear responses change when they are exposed to human visual cues alone or to additional cues, such as human voices and pet visual and acoustic cues. We used alert distance (AD) and flight initiation distance (FID) as quantitative indicators of fear. AD and FID measure the distance at which prey begins vigilant behaviour and initiates flight/escape behaviour in response to an approaching predator, respectively (Dumont et al., 2012; Stankowich, 2008; Uchida et al., 2019). Both measures index fear but capture different decision processes (Dumont et al., 2012; Petrelli et al., 2017; Stankowich, 2008; Uchida et al., 2019). AD measures vigilance, which can begin once a potential predation risk is detected, even if the immediate threat is uncertain (Cooper & Blumstein, 2014; Uchida et al., 2019). FID measures flight or escape, which imposes substantial costs (e.g., energy expenditure) and requires evaluating the urgency of the risk and balancing the costs and benefits of fleeing (Ydenberg & Dill, 1986; Uchida et al., 2019). These differences suggest that acoustic and visual cues may differentially influence early detection (AD) and escape decisions (FID). For example, AD may increase when predators produce acoustic cues detectable from relatively long distances, whereas FID may increase when prey receives visual predator cues, because visual information can more reliably signal predator presence (Arteaga- Torres et al., 2020; Mathot et al., 2024).

In this study, we measured AD and FID in Yezo sika deer (*Cervus nippon yesoensis* Heude) in open landscapes in Hokkaido, northern Japan, during both daytime and nighttime trials. We presented human visual cues alone and in combination with additional cues–human acoustic cues, dog visual cues, and dog acoustic cues–in multiple cue combinations. We focused on dogs as representative pets for two reasons. First, among predators that humans have domesticated (e.g., cats, hawks, eagles, owls) (Driscoll et al., 2009; Farner et al., 2013), dogs are the most common pets worldwide, with an estimated global population of approximately one billion (Bryce, 2021; Gompper, 2014; Vanak & Gompper, 2009). Second, dogs are frequently used as experimental stimuli in studies of fear effects, and they may act as potential predators of deer because wolves (the same species as domestic dogs) prey on deer (Bryce, 2021; McGann et al., 2024). We also selected Yezo sika deer as the focal prey species for two reasons. First, their high abundance in Hokkaido makes them readily detectable, allowing us to measure AD and FID efficiently in open landscapes. Second, Yezo sika deer belong to the cervids that cause agricultural damage (Elisa et al., 2016; Hata et al., 2021; Lande et al., 2014; Trdan & Vidrih, 2007; Woodroffe et al., 2005), and rising cervid populations worldwide pose a major challenge for wildlife management (Conover, 1998; Takatsuki, 2009; Warren, 2011). By identifying the cues that drive deer fear responses to humans and dogs, our study may help inform management strategies aimed at reducing crop damage and zoonotic disease risk by influencing deer movement and space use. We tested the following two hypotheses (Fig.1).

1. AD and FID would increase when the human acoustic cue, the dog visual cue, and the dog acoustic cue are added to the human visual cue.
2. AD would be more strongly influenced by the addition of acoustic cues, whereas FID would be more strongly influenced by the addition of visual cues.

**Figure. 1.**
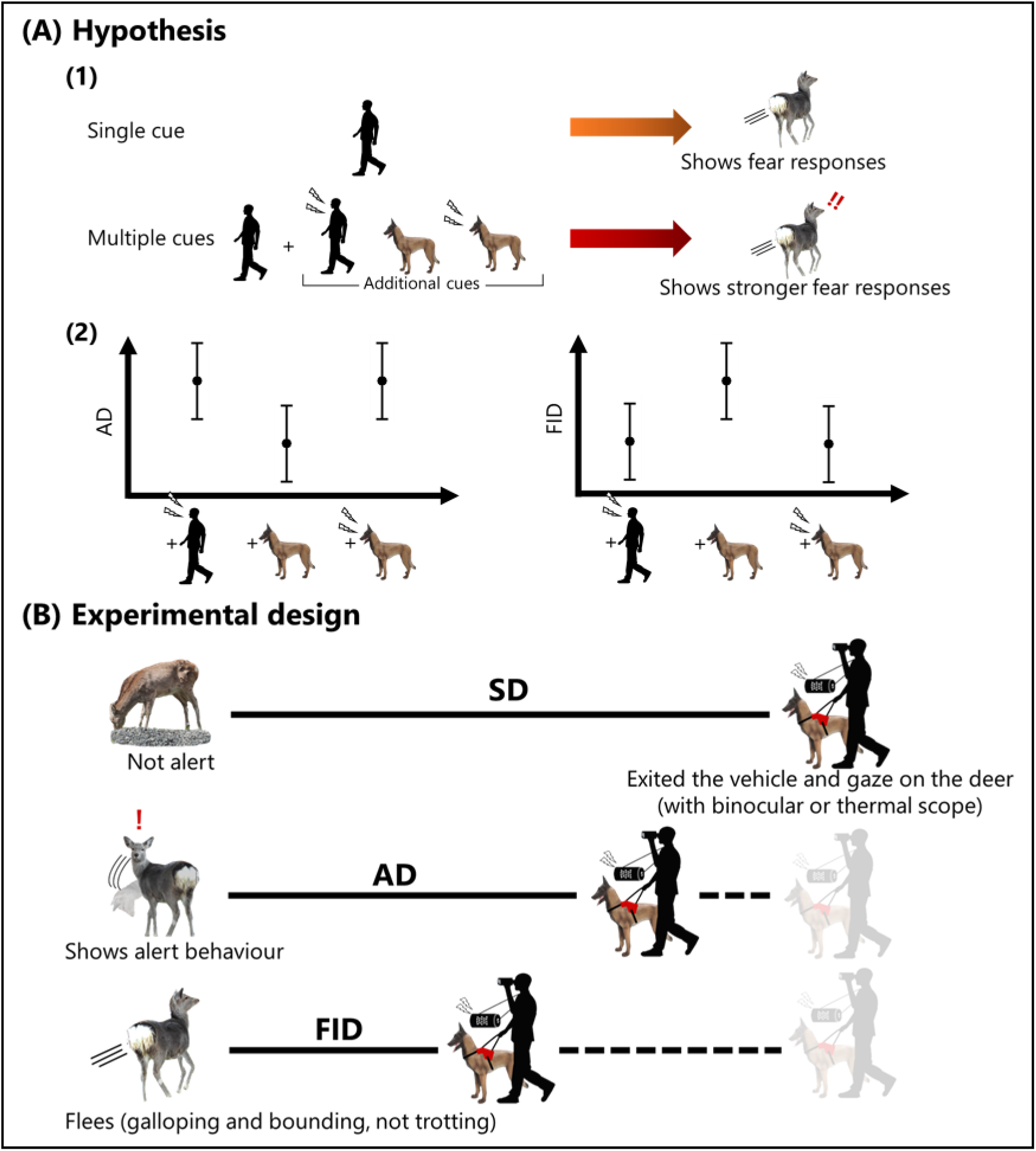
**(A)** Expected behavioural response of sika deer to single and multiple cues (1) and expected AD and FID adding each human and pet cues to the human visual cue (2). **(B)** Design of the FID and AD measurement. A surveyor approached focal individual(s) and recorded starting distance (SD), AD, and FID.

## Methods

### The Target Species

Yezo sika deer, the focal species of this study, are a subspecies of sika deer (*Cervus nippon* Temmink). It is a large herbivorous ungulate that commonly uses forest edges but also occupies a wide range of habitats. In Hokkaido, Yezo sika deer occur across most areas except in areas with high human population density. Migratory females had the largest home ranges (21.1 ± 5.5 km²) among the population in Kushiro-shitsugen National Park, Hokkaido (Takafumi et al., 2017). The rut occurs in autumn, during which males form harems (Ohdachi et al., 2015). Outside the rut, individuals may form single-sex groups (Ohdachi et al., 2015). Yezo sika deer are mainly active at night but often remain active during the day. Although they represent an important game species in Japan, increasing populations have become a serious social issue due to crop and forestry damage and collisions with vehicles and trains (Ohdachi et al., 2015).

### Cue Preparation

To measure AD and FID in our focal species, we used a human visual cue and combinations of human acoustic and dog visual and acoustic cue(s). A single surveyor (first author, male) conducted all measurement trials, and the human visual cue was kept constant across trials. We used a fibre-reinforced plastic Belgian Shepherd dog decoy (height: 88 cm; body length: 90 cm) as the dog visual cue (Fig.1), as its size is representative of a potential predator of the focal species. We used 50 unique male conversation audio files from the listening section of the Japanese-Language Proficiency Test (https://www.jlpt.jp/about/index.html). Using audio-editing software (Audacity, The Audacity Team), we segmented those files into 5s clips (sampling rate: 44.1 kHz), avoiding unnatural truncation of speech. We finally obtained 232 playback files as the human acoustic cues. We also used dog barking audio files from freely available online sources (https://mixkit.co/free-sound-effects/dog/; https://pixabay.com/sound-effects/search/dog%20barking/). We then extracted 1s single barks and repeated them five times to create 5-s playback sequences (one bark per second). We finally created 304 playback files as the dog acoustic cues. We also created a 5-s white-noise playback file using a freely available online source (https://soundqualitylab.com/test_tones/02_noise/01_whitenoise/) to confirm that deer did not respond simply to sound per se (Table 1). Finally, to confirm that deer recognized the dog decoy as an approaching dog, we also used the same decoy covered with a black plastic sheet (90 cm × 100 cm). We finally comprised 10 variations of visual and acoustic cues (Table 1).

**Table 1.** Cue combinations and the number of trials used to measure AD and FID.

| Human<br>visual cue<br>(surveyor<br>approach) | Human<br>acoustic cue<br>(voice) | Dog visual<br>cue (decoy) | Dog<br>acoustic cue<br>(barking) | Covered<br>dog decoy<br>(control) | White<br>noise<br>(control) | No. of trials<br>(FID ≤ 150 m) | No. of excluded<br>trials |
| --- | --- | --- | --- | --- | --- | --- | --- |
| + | - | - | - | - | - | 33 | 1 |
| + | + | - | - | - | - | 30 | 4 |
| + | - | - | + | - | - | 28 | 5 |
| + | - | - | - | - | + | 32 | 2 |
| + | - | + | - | - | - | 33 | 1 |
| + | + | + | - | - | - | 30 | 4 |
| + | - | + | + | - | - | 29 | 5 |
| + | - | + | - | - | + | 31 | 0 |
| + | - | - | - | + | - | 10 | 0 |
| + | - | - | + | + | - | 10 | 0 |

“+” indicates cues used in each trial, and “-” indicates cues not used.

For each playback sound file, we calibrated the amplitude in a closed soundproof room. In this calibration, we ensured that each playback sound had 80 ± 2 dB A-weighted equivalent continuous sound level (LAeq) at 1 m from the speaker (CHARGE 3, JBL, California, USA).

### Study Area and Survey Period

We conducted field surveys across an area of approximately 800 km² in the Iburi, Hidaka, and Ishikari regions of Hokkaido, northern Japan, including the Tomakomai Experimental Forest (Fig.2). The study sites consisted of open, high-visibility habitats such as farmland and abandoned land, as well as urban areas. We conducted measurement trials at flat, unobstructed sites with low vegetation, where deer could clearly see the entire dog decoy. We carried out all trials on days with favourable weather (avoiding strong winds and heavy rain) from 10 August 2024 to 29 May 2025.

**Figure. 2.**
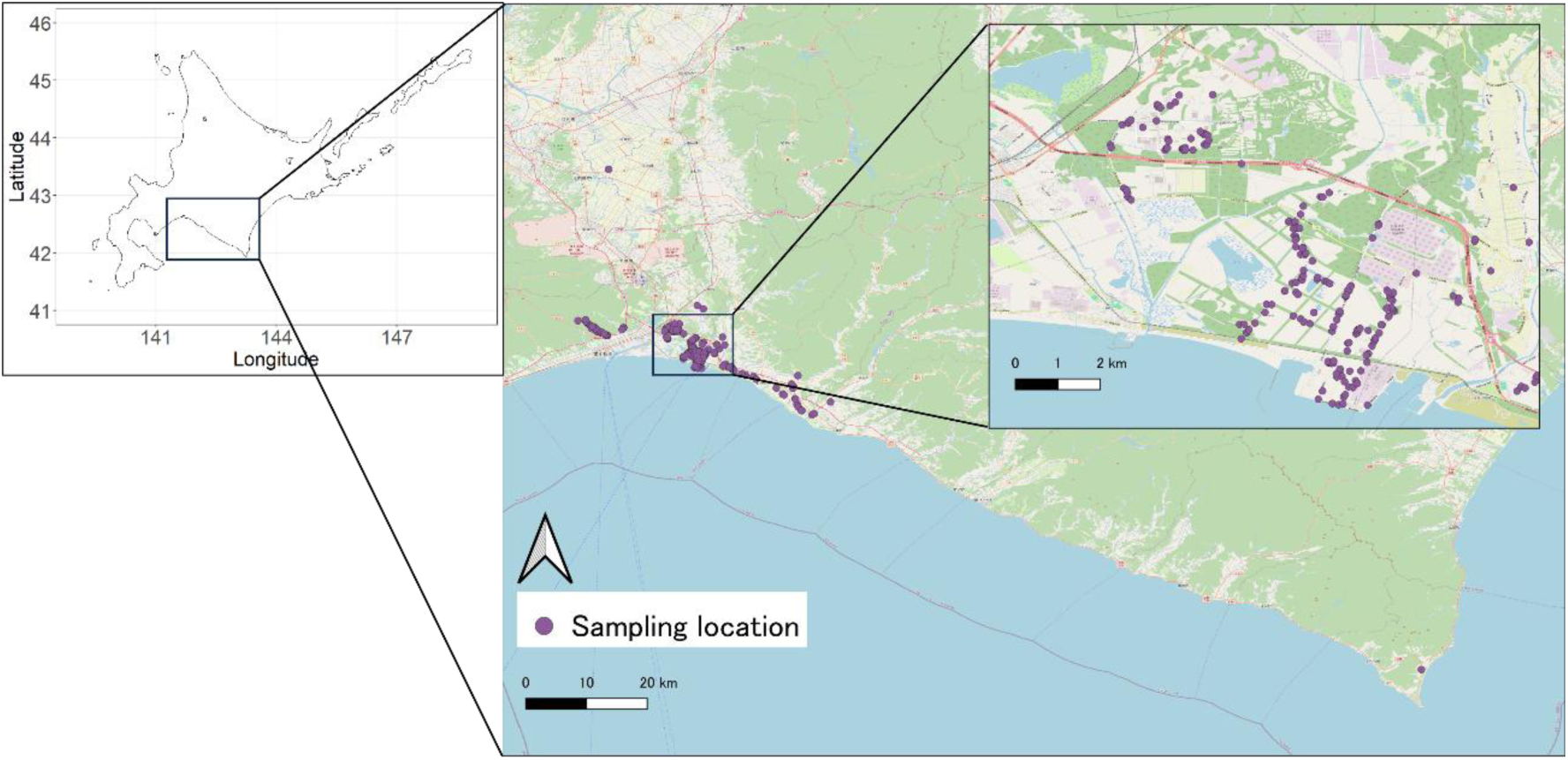
Map of Hokkaido (left) and the study area (right; 42.0-43.0°N, 141.6-143.2°E), showing the locations where AD and FID were measured after excluding trials with FID > 150 m. Basemap: OpenStreetMap contributors (https://www.openstreetmap.org/copyright/ja).

### The Design of Measurement Trials

We searched for deer from a vehicle or motorcycle at locations of suitable habitats. We tried to detect deer at the greatest possible distance using a binocular (MONARCH 7, 8×42, Nikon, Tokyo, Japan) during the day (from sunrise to sunset) or with thermal scopes (Thermal Imaging Monocular EYE III Series EL35, InfiRay, Yantai, China; CONDOR CQ35L, HIKMICRO, Hangzhou, China) at night (from sunset to sunrise). We selected deer showing no obvious signs of vigilance (i.e., not looking directly at us) as the targets (Supplementary Video S1 and S2). Because most deer in the study area inhabit areas frequently used by humans and dogs, we assumed that humans and dogs were not novel objects for deer, except for immature individuals (< 1-year). We therefore excluded these immature individuals from the measurement trials.

Once a focal deer was located for measurement, the surveyor exited the motorcycle or vehicle and, when necessary, set up the dog decoy before initiating a trial. When a group of deer was detected, the surveyor focused on the closest individual. Previous studies have shown that the surveyor’s clothing colour and approach speed can influence AD and FID (Golawska et al., 2024; Kosako et al., 2008). Accordingly, the surveyor wore the same clothing in all trials (orange outer layer and beige trousers) and approached the deer along a straight line at a constant speed (< 1m/s). For a trial with the dog visual cue, the surveyor carried the decoy slightly above the ground using a harness attached to it, thereby mimicking a person walking a dog. For a trial with an acoustic cue (human voice, dog barking, or white noise), the surveyor selected the assigned playback file on a portable player (WALKMAN, Sony, Tokyo, Japan) and broadcast it repeatedly in loop mode from a speaker (CHARGE 3) worn around the neck, from the initiation of approach to the end of the measurement trial. To avoid pseudo-replication, we used a different playback file in each trial (except for white noise), following a pre-assigned order (Whitehouse et al., 2024). If a vehicle passed during a trial, we paused and excluded the trial from the analyses, because we could not determine whether the deer responded to the vehicle or to the surveyor (this occurred once).

During each trial, we recorded AD and FID. We defined FID as the linear distance between the surveyor and the location of the deer that initiated escape behaviour (i.e., typically running away from the surveyor). We defined AD as the linear distance between the surveyor and the location of the deer that first became vigilant (i.e., began gazing at the surveyor). To calculate AD, the surveyor marked the position at the onset of vigilance (e.g., with a peg), measured the distance moved from that point to the position at flight initiation, and added this distance to FID. We also measured the linear distance between the surveyor and the initial detection location of the focal individual (hereafter referred to as starting distance [SD]). For this calculation, the surveyor marked the position at detection, measured the distance from that point to the position at flight initiation, and added this distance to FID. These measurements were conducted with laser rangefinders (ULT-X 800, TecTecTec!, France; CONDOR CQ35L).

### Ethical Note

This study involved behavioural observations of free-ranging Yezo sika deer and did not involve any invasive procedures. After each measurement trials, the surveyor immediately ceased the approach, left the area, and did not pursue the animals further. To avoid using a live dog in the experiments, we used a life-sized dog decoy as a welfare-conscious alternative. Playback sounds were presented only during the brief period required for the standardized approach and ceased immediately after the trial, thereby avoiding prolonged exposure.

We conducted all trials in accordance with the ASAB/ABS Guidelines for the treatment of animals in behavioural research and teaching (https://doi.org/10.1016/j.anbehav.2025.123421). We followed legal requirements according to the Japanese Wildlife Protection and Hunting Law L.N. 88. Fieldwork was approved by authorization decrees F2024122B of 26/July/2024 and F2025071B of 9/May/2025 from Tomakomai Experimental Forests. We did not seek specific ethical committee approval as this was not required for noninvasive wildlife research.

### Measurement of Environmental Variables

We recorded ambient light level, background noise level, deer group size, mean wind speed, maximum wind speed, season (rutting vs non-rutting), and time of day as environmental variables that could influence AD and FID. The ambient light level may affect the distance at which the focal deer can visually detect the surveyor (Stankowich, 2008). We measured illuminance (lx) at the flight initiation location with an illuminance UV recorder (TR-74Ui, T&D, Tokyo, Japan). We pointed the sensor upwards, recorded three measurements to two decimal places at 10-s intervals, and then calculated their mean. The average ambient light level was 4856 ± 15724 lx (mean ± SE). The background noise level may affect the audible range of the playback (Darras et al., 2016). We recorded 1-min LAeq to one decimal place with a sound pressure meter (TYPE6236, ACO, Miyazaki, Japan) at the same location used for the illuminance measurements (measurement range: 28-130 dB). The average background noise level was 40.7 ± 6.5 dBA. Group size may influence the focal deer’s response through collective behaviour, and such effects may be stronger in larger groups (Aastrup, 2000; de Boer et al., 2004; Stankowich, 2008). We recorded group size as the number of individuals that fled simultaneously with the focal deer. The average group size was 7 ± 10 individuals. Wind speed may influence the audible range of playback and the detectability of olfactory cues from the surveyor. Using a digital anemometer (MT-EN1A, Mother Tool, Nagano, Japan), we recorded the mean and maximum wind speed over 30 s to one decimal place. The average mean and maximum wind speeds were 0.4 ± 0.9 m/s and 0.6 ± 1.2 m/s, respectively. Seasonal behavioural shifts may influence AD and FID (Ciuti et al., 2008). We classified trials as rutting season (October-November; N = 137) or non-rutting season (December-September; N = 129). Time of day or diel changes in deer behaviour and visibility may influence AD and FID. We classified trials as daytime (sunrise-sunset; N = 86) or night-time (sunset-sunrise; N = 180) because the same clock time can correspond to different times of day across seasons.

To minimize the possibility that deer experienced playback sounds from another measurement trial, we spaced each trial by at least 150 m, based on the audible range of the playback (See Supporting Information; Fig.S1). To reduce habituation, we conducted repeated trials at the same location only after an interval of at least one week. We excluded two trials that did not meet these criteria. The same surveyor (first author) conducted all trials.

### Statistical Analyses

We fitted two linear mixed-effects models (LMMs), one each for AD and FID. We compared models fitted to the raw responses with those fitted to log-transformed responses, and selected the raw response models because of the higher R² values (See Supporting Information; Appendix S1 and Table S1). Because the playback sounds (human voice, dog barking, and white noise) attenuated with distance and reached 32.5 ± 2.5 dBA at 150 m from the speaker (See Supporting Information; Fig.S1), deer might be unable to detect the playback sounds under high background-noise conditions. We therefore restricted the analyses to trials with FID ≤ 150 m.

Each model included the following five binary explanatory variables: (1) dog decoy, (2) covered dog decoy, (3) human voice, (4) dog barking, and (5) white noise. We included a human visual cue (the approaching surveyor) in all trials, and occasionally combined it with one or two additional cues, resulting in ten cue-combination patterns (Table 1). We therefore treated the presence of the human visual cue as the baseline condition (i.e., intercept) of each model. We also included SD, ambient light level, group size, mean wind speed, and season (rutting vs non-rutting) as additional explanatory variables. We excluded maximum wind speed and time of day because of high correlations (|r| >0.6) with mean wind speed and ambient light level, respectively. We also excluded the background noise level because it showed high multicollinearity (VIF > 5). Because deer have wide home ranges and individuals may move among measurement points, we may have measured the same individual more than once. Based on the maximum home-range size (21.1 ± 5.5km^2^; Takafumi et al., 2017), we overlaid a 5 km × 5km grid and assigned the same location ID to measurement points that fell within the same grid cell. We included location ID as a random effect in the LMMs. To account for the effect of differences in the lengths of voice segments, we quantified the proportion of “sound-present” segments in each file with a -35 dB peak-level threshold and a minimum silence duration of 0.010 s. Although the voice segment proportion ranged from 45% to 81%, a preliminary analysis showed that its effect on deer behaviour was not significant (FID: β=-6.00, p = 0.374; AD: β= -12.1, p = 0.197; see Supporting Information Appendix S2 and Table S2 for details)

To compare effect sizes in each model, we calculated 95% confidence intervals (CIs) for the estimated coefficients and considered effects statistically significant when the 95% CI did not include zero. We also calculated model-based estimated values of AD and FID. We fitted all models in R version 4.4.1 using the ‘lme4’ package (Bates et al. 2014; R Core Team, 2024). We assessed multicollinearity using the ‘car’ package (Fox & Weisberg, 2019), calculated 95% CIs using the ‘MASS’ package (Venables & Ripley, 2002), calculated the R² values using the ‘performance’ package (Lüdecke et al., 2021), and obtained estimated marginal means of AD and FID using the ‘emmeans’ package (Lenth, 2024).

## Results

### Field Measurement Trials

Over a total of 48 survey days, we collected 290 AD/FID observations of Yezo sika deer across 24 location IDs (location IDs 1-24; Fig. 2; see Supporting Information Fig.S4 for details). We analysed 266 observations after excluding trials that did not meet our criteria for the analyses (> 150 m between trials, ≥ 1-week interval between repeat trials at the same site, and FID ≤ 150 m; Table 1). After exclusion, the sampling period and the total number of survey days remained unchanged, but the dataset covered 22 location IDs.

### Alert Distance

AD was estimated at 80.8 ± 4.74 m (estimate ±SE, 95% CI = [71.3, 90.3]) in trials with only the human visual cue. The addition of dog barking and white noise increased AD by 18.4 ± 6.46 m (estimate ± SE, ≈23%; 95% CI = [5.71, 30.6]; Table 2, Fig. 3) and by 17.0 ± 5.26 m (estimate ± SE, ≈21%; 95% CI = [6.65, 26.9]; Table 2, Fig.3), respectively. In contrast, the addition of the human voice, the dog decoy, or the covered dog decoy did not affect AD (Table 2). No significant interactions were detected between the dog decoy and either dog barking or human voice (Table 2). Together, these results suggest that deer relied more on acoustic than visual cues when initiating vigilance, and that cue effects on AD were additive. We also found that AD increased with SD (β ± SE = 0.523 ± 0.0283, 95% CI = [0.467, 0.583]) and was higher in the non-rutting season (β ± SE = 9.19 ± 4.11, 95% CI = [1.26, 17.0]; Table 2).

**Figure. 3.**
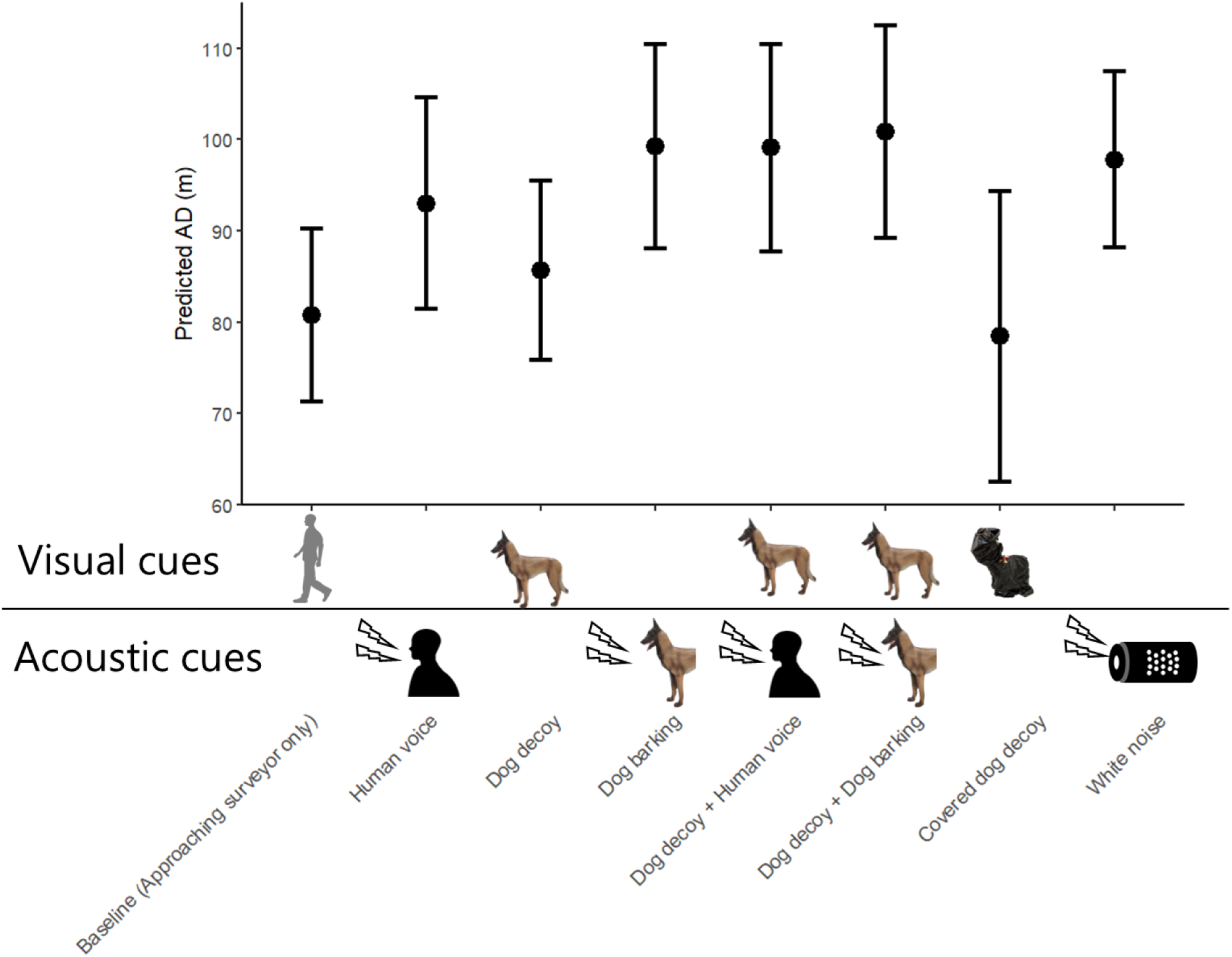
Model-based estimated alert distance (AD) for each cue added to the human visual cue. Black points and error bars indicate the estimated values and the 95% confidence intervals, respectively. N =266.

**Table 2.**
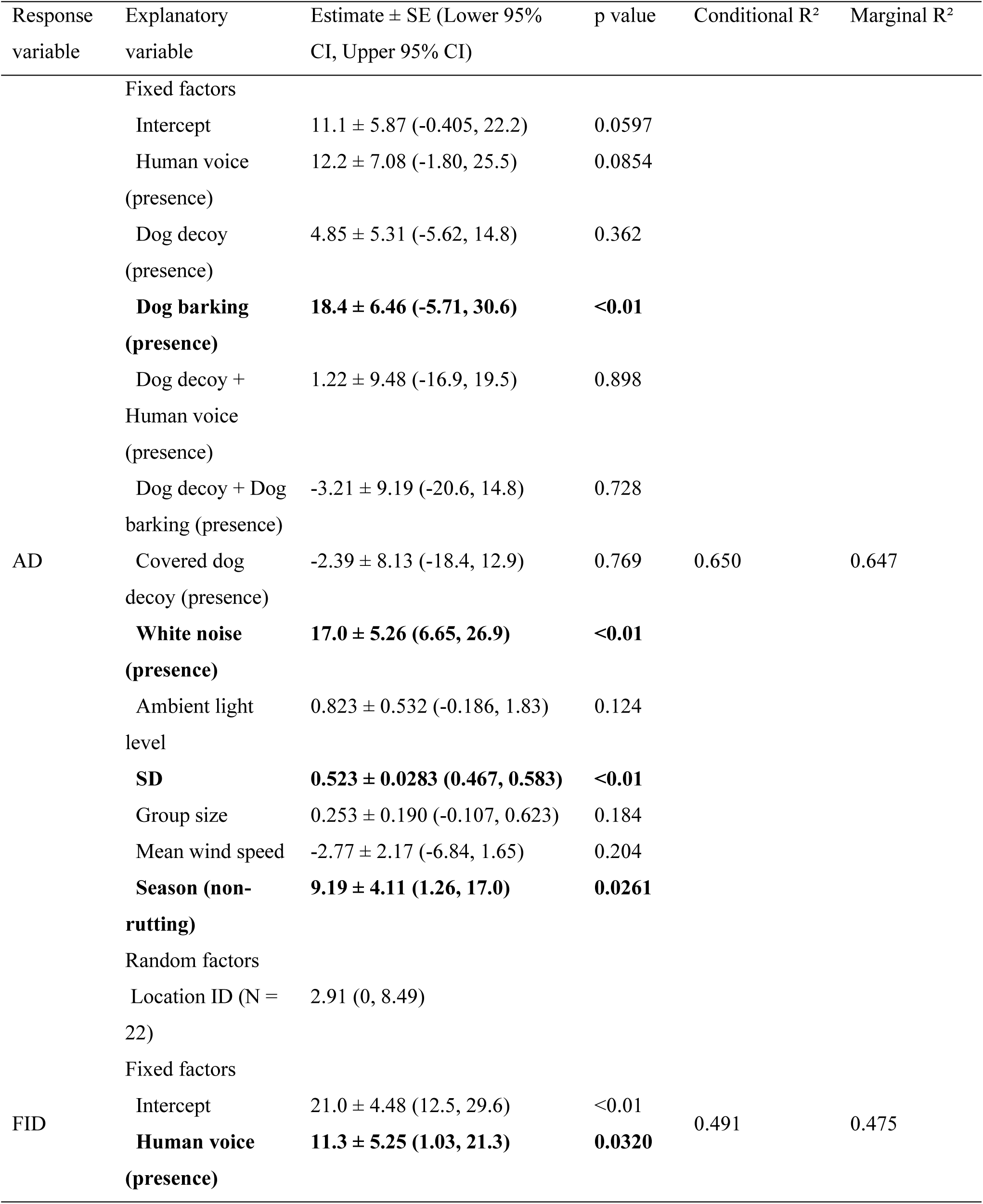
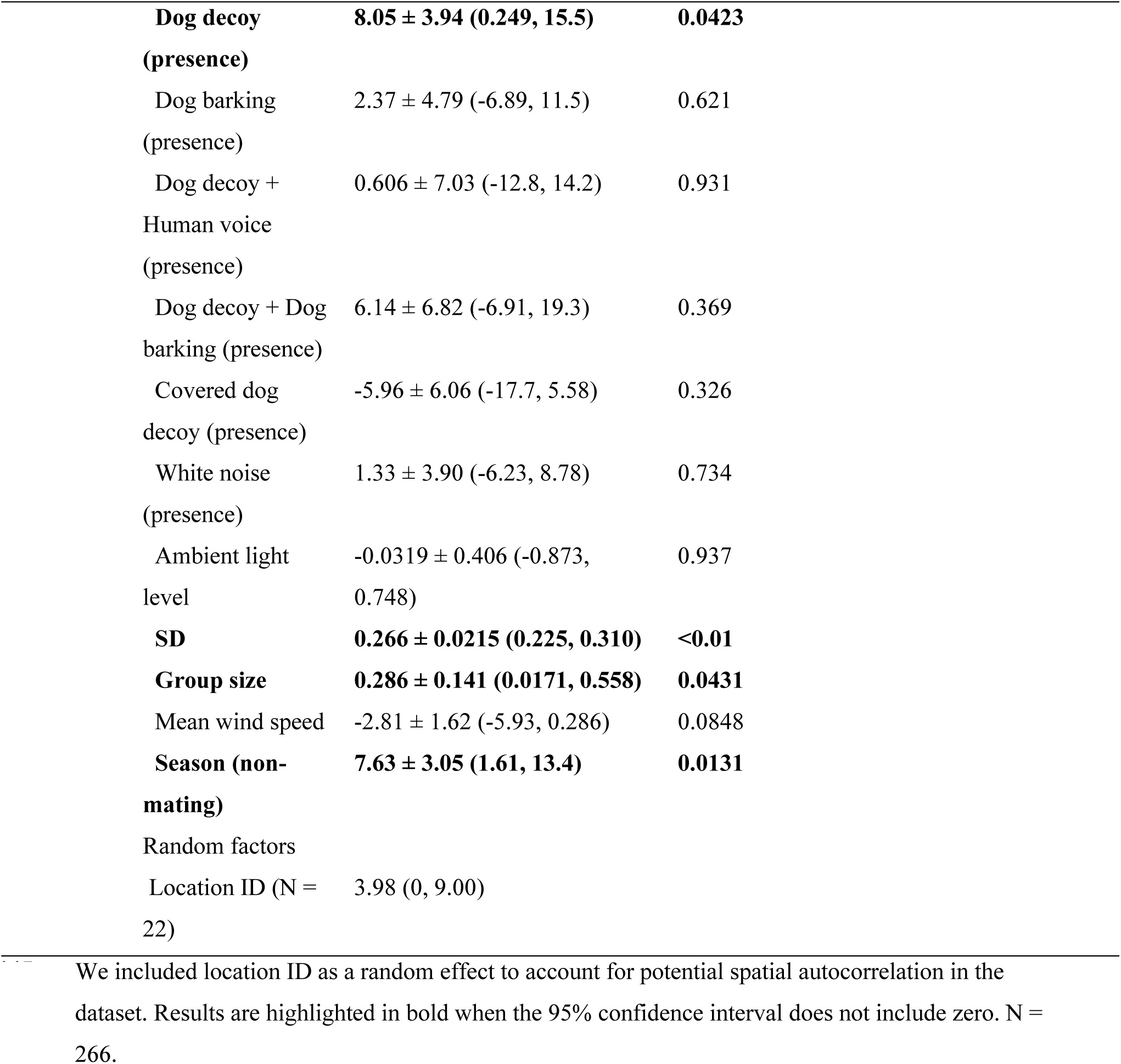
Results of linear mixed-effects models (LMMs) evaluating the effects of cues and environmental factors on alert distance (AD) and flight initiation distance (FID).

### Flight Initiation Distance

FID was estimated at 57.1 ± 3.69 m (estimate ± SE, 95% CI = [49.7, 64.5]) in trials with only the human visual cue. The addition of human voice and the dog decoy increased FID by 11.3 ± 5.25 m (estimate ± SE, ≈20%; 95% CI = [1.03, 21.3]; Table 2, Fig.4) and by 8.05 ± 3.94 m (estimate ± SE, ≈14%; 95% CI = [0.249, 15.5]; Table 2, Fig.4), respectively, suggesting that deer initiated escape behaviour at greater distances when they perceived human acoustic cues as well as the dog visual cue in addition to the human visual cue. Moreover, the addition of the covered dog decoy did not affect FID (Table 2), suggesting that deer likely responded to the dog-like visual features of the decoy.

**Figure. 4.**
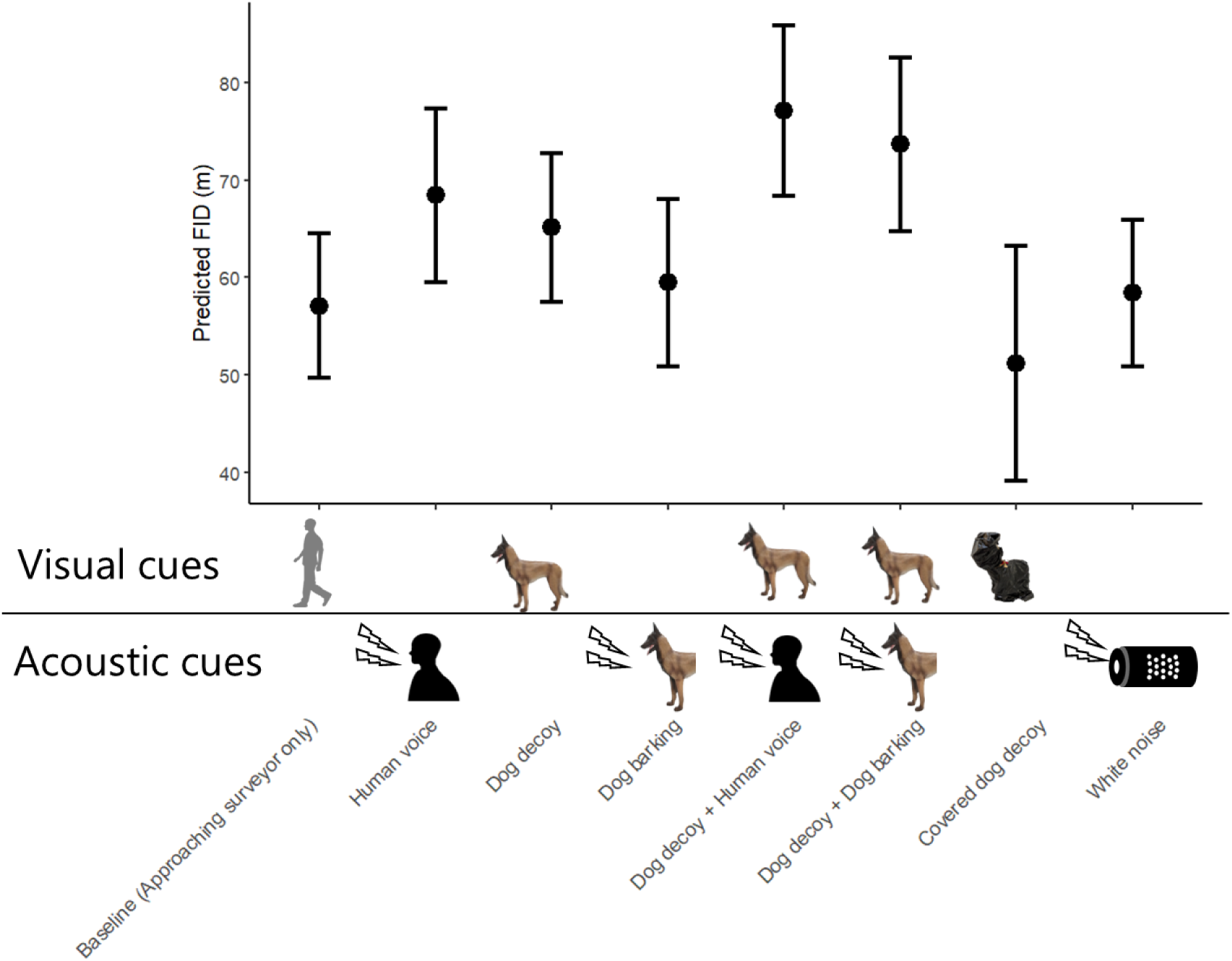
Model-based estimated flight initiation distance (FID) for each cue added to the human visual cue. Black points and error bars indicate the estimated values and the 95% confidence intervals, respectively. N =266.

In contrast to the longer FID observed in deer exposed to the dog visual cue, the addition of dog barking did not affect FID (Table 2), suggesting that deer relied more on visual than acoustic cues when responding to dog-related risk. No significant interactions were detected between the dog decoy and either dog barking or human voice (Table 2). Furthermore, white noise did not affect FID (Table 2), supporting the interpretation that deer did not exhibit their escape response solely in response to sound playback. We also found that FID increased with SD (β ± SE = 0.266 ± 0.0215, 95% CI = [0.225, 0.310]) and group size (β ± SE = 0.286 ± 0.141, 95% CI = [0.0171, 0.558]), and was higher in the non-rutting season (β ± SE = 7.63 ± 3.05, 95% CI = [1.61, 13.4]; Table 2).

## Discussion

It remains unclear how wildlife fear responses to humans vary with the presence of pets, as well as with the types and combinations of cues perceived from humans and pets. We addressed this gap by examining whether human acoustic cues increased wildlife fear responses to human visual cues, as measured by both alert distance (AD) and flight initiation distance (FID). We also tested whether dog-related cues further amplified these fear responses beyond those elicited by human visual cues alone. Consistent with Hypothesis 1, both AD and FID increased when additional cues were added to the human visual cue, suggesting that talking and walking with dogs may intensify perceived risk for wildlife. We also found cue-specific differences between response stages. AD was more strongly influenced by additional acoustic cues, whereas FID was more strongly influenced by both additional visual and acoustic cues, partially supporting Hypothesis 2.

### Similarities and Differences Between Human and Predator Cue Effects

Our findings align with previous studies demonstrating that multimodal cues from natural predators increase prey fear responses (Jones et al., 2024; Mukherjee et al., 2024; Munoz & Blumstein, 2012; Sih, 1986), and extend this framework to fear responses towards humans. On the other hand, few studies have explicitly tested the incremental effect of adding direct predator acoustic cues to visual cues on fear responses. To our knowledge, only Lukas et al. (2021) has tested how adding bird acoustic cues to bird visual cues affects fish escape behaviour. However, fish escape options were constrained (e.g., the maximum dive depth) because that study measured responses under laboratory conditions. Our field experiment likely provides a more realistic assessment of the effect of adding acoustic cues on escape behaviour, where escape distance was not physically constrained. Moreover, group size was a significant factor in the full FID model (Table 2), but it was not in the human-unimodal cue subset, and reappeared in the human-multimodal cue subset (See supporting Information Table S3 for details). This result is consistent with evidence that multimodal cues can yield more robust responses, whereas responses to unimodal cues often show greater variability (Mathot et al., 2024). Field experiments testing the combined effects of multiple sensory cues may therefore provide more robust inferences about animal fear responses, highlighting the importance of considering multiple sensory modalities in natural settings.

Our research also demonstrated that walking with pets—a human-specific activity pattern not shared by other predators—increases wildlife fear responses. In natural conditions, prey commonly face risk from multiple predator species, but predators typically occur independently in space and time (Sih et al., 1998). In contrast, humans and pets, particularly dogs, often move together, which can present human- and pet-cues simultaneously and consistently to wildlife. Walking with pets may therefore create situations analogous to the co-occurrence of two predator species, potentially at higher frequencies than in natural conditions. Previous studies, however, largely overlooked fear of pets, as they relied on tools such as camera traps rather than direct behavioural measurements and often used simplified experimental designs, such as playback of human vocalizations. Our results therefore emphasize the importance of incorporating realistic human activity patterns, such as walking with dogs, into behavioural measurements of response to human disturbance to improve ecological realism (e.g., Zeller et al., 2024).

### Differences in Cues Affecting AD and FID

Previous research has reported that vigilance can begin when wildlife detects a predation risk, while escape requires evaluating the urgency of the risk (Cooper & Blumstein, 2014; Uchida et al., 2019; Ydenberg & Dill, 1986). We therefore hypothesized that AD would be increased when prey perceived acoustic cues that reach farther, whereas FID would be increased when they received visual predator cues that substantiate predator approach. Consistent with our prediction, dog acoustic cues increased AD. However, human acoustic cues, as well as the dog visual cue, increased FID. This result was contrary to our prediction and to previous evidence that visual cues of predation risk outweigh acoustic cues (Arteaga-Torres et al., 2020). One possible explanation for this discrepancy is that animals adjust the relative importance of cues depending on predator species, and that acoustic cues become more important than visual cues when assessing risk from humans. Human voices contain distinctive acoustic signatures, including characteristic resonances produced in the vocal tract, that differ from those of other animals (Fitch, 2000). Animals can perceive those resonances with an accuracy rivalling that of humans (Owren, 1990; Sommers et al., 1992). Human voices may therefore provide information that helps animals identify an approaching threat as human (Fitch, 2000). Future research could evaluate whether prey adjust the relative importance of visual, acoustic, and other cues across predator species, and whether distinctive features of human voices enable prey to identify approaching threats as human and thereby increase reliance on acoustic cues during risk assessment.

### Limitations and Future Research Directions

Three limitations should be considered in future work. First, although the sex of the targeted individual can influence FID (Morelli et al., 2025; Stankowich, 2008), our models were unable to include it because determining deer sex at night was technically difficult. Evidence from other artiodactyls suggests that sex-related differences in FID can be substantial (e.g., ∼18m in European bison (*Bison bonasus* Linnaeus; Haidt et al., 2018), potentially comparable to the cue effects observed in this study (e.g., adding a human voice: + 11.3 m; adding the dog barking: + 8.05 m). However, conclusions regarding sex differences in FID remain inconsistent across species (Stankowich, 2008), highlighting the need for evaluation in our target species. Second, we restricted our analyses to FID ≤ 150 m because of the limited audible range of playback sounds, and this may limit the generality of our findings to more sensitive individuals or populations (FID > 150 m). How fear responses change when acoustic cues remain detectable beyond this distance remains unclear and warrants future investigation. Finally, we did not explicitly account for spatial or individual variation in habituation to humans. Birds, mammals, and reptiles show shorter FIDs in human-habituated populations (Samia et al., 2015). Repeated human approaches can also alter FID through habituation or sensitisation (Mohring et al., 2025; Runyan & Blumstein, 2004). In our models, however, the variance explained by random effects was small, suggesting limited among-location variation in AD and FID. Although individual-level variation in habituation to humans may still matter, differences in habituation among areas likely had only a minor influence on our main conclusions.

### Management Implications

Our results underscore the importance of considering both human and dog cues in conservation and management. Whether to ban walking with pets, such as dogs, in sensitive areas remains controversial (Banks & Bryant, 2007). Our findings provide partial support for the effectiveness of such restrictions, given that reducing human- and dog-related cues may help mitigate wildlife disturbance. Fleeing—associated with higher energetic and opportunity costs than vigilance (Lima & Dill, 1990)—typically occurs only after initial detection if humans continue to approach, and in our study was primarily influenced by human acoustic cues and dog visual cues, both of which are relatively tractable targets for management through signage, visitor guidance, and spatial zoning (e.g., restricted areas). Measures, such as keeping dogs on leads, discouraging close approaches to wildlife, encouraging quiet behaviour, and designating pet-free areas, may therefore help reduce high-cost escape responses while allowing continued recreational use. In contrast, dog barking, which affected deer vigilance responses, may be more difficult to control directly. To minimise vigilance-related disturbance, practical management actions could include maintaining suitable distances between visitors with dogs to reduce dog-dog interactions that can trigger barking, alongside clear guidance on dog-handling practices in natural areas. Nevertheless, managing how humans and dogs behave—rather than simply restricting their presence—may offer an effective and practical approach to reducing wildlife disturbance.

## Data availability

Data available from the Github repository (https://anonymous.4open.science/r/Multimodal_Human_Pet_Cue_Fear/README.md)

## Acknowledgements

We thank the staff in the Tomakomai Experimental Forest for supporting our field survey. We also thank Senzaki Lab members for providing useful comments for the improvement of the methodology and manuscript. Anonymous reviewers provided useful comments for the improvement of the manuscript. This study was supported by the Japan Society for the Promotion of Science KAKENHI (23K26936).

