## Supplementary materials for "Multimodal human and pet cues intensify wildlife fear responses"

For the manuscript

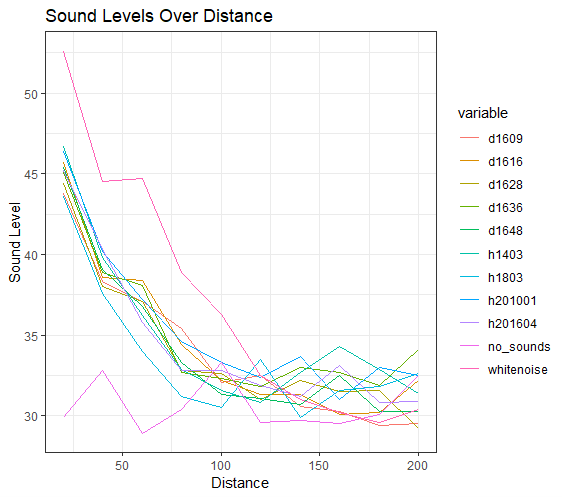

**Figure. S1.** Attenuation of playback sound levels with increasing distance from the speaker. Dog-barking playbacks are d1609, d1616, d1628, d1636, and d1648; human-voice playbacks are h1403, h1803, h201001, and h201604; White-noise playback is *whitenoise*. *no_sounds* indicates measurements of environmental noise only (no playback). The x-axis shows distance from the speaker (m), and the y-axis shows the equivalent continuous A-weighted sound pressure level over 1 min (LAeq,1min; dBA).

We conducted attenuation measurements along a straight forest road in the Tomakomai Experimental Forest, Hokkaido, northern Japan. We mounted the speaker on a tripod at a height of 1.2 m. Using a laser rangefinder, we marked measurement points at 20-m intervals from 20 m to 200 m (20, 40, 60, 80, 100, 120, 140, 160, 180, and 200 m) and recorded sound levels at each point with a sound level meter. Even for white noise, which showed the least attenuation among the tested playbacks, sound levels became comparable to ambient environmental noise at approximately 150 m.

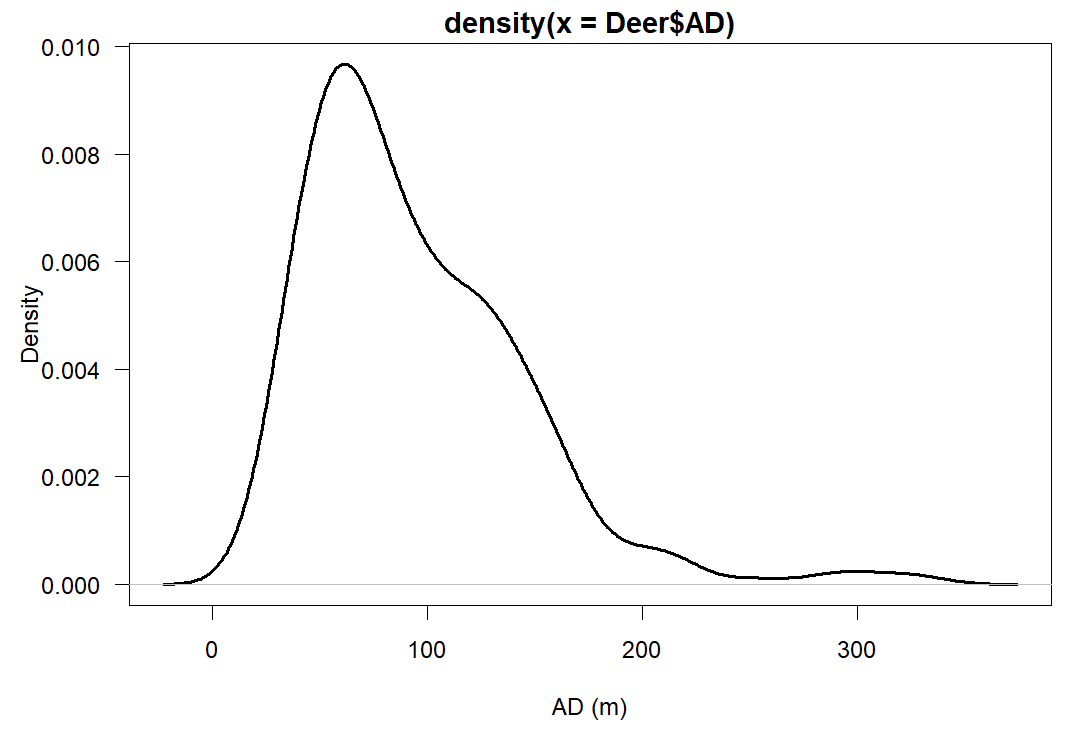

**Figure. S2.** Kernel density estimate of the distribution of observed alert distance (AD).

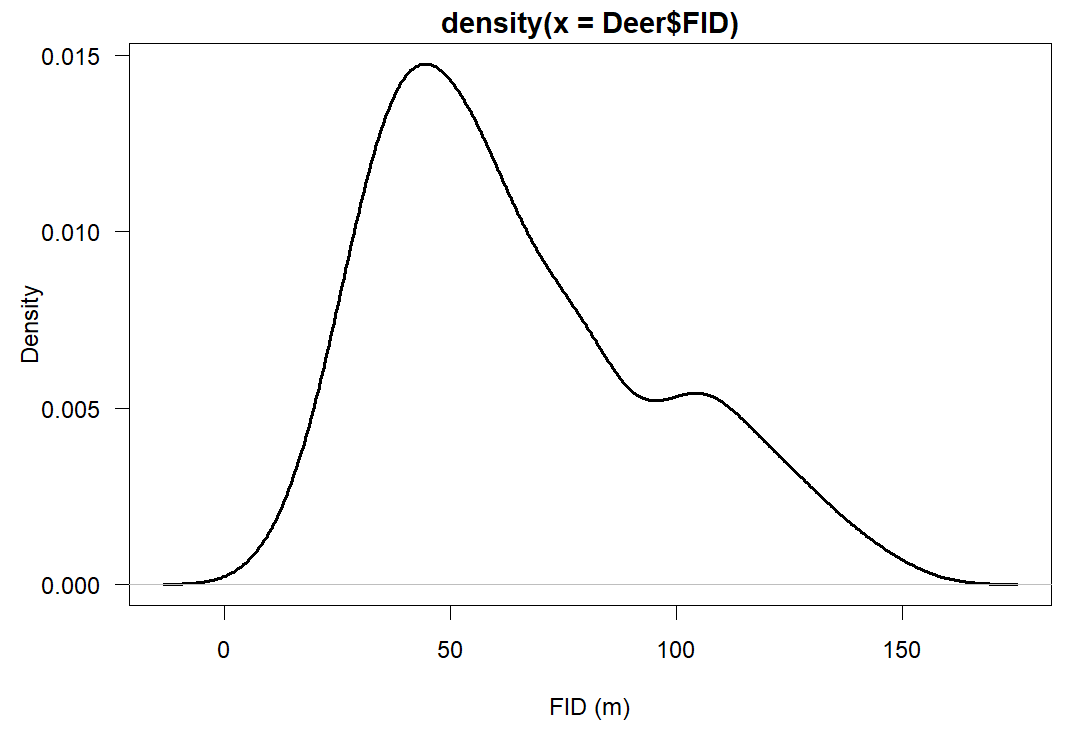

**Figure. S3.** Kernel density estimate of the distribution of observed flight initiation distance (FID).

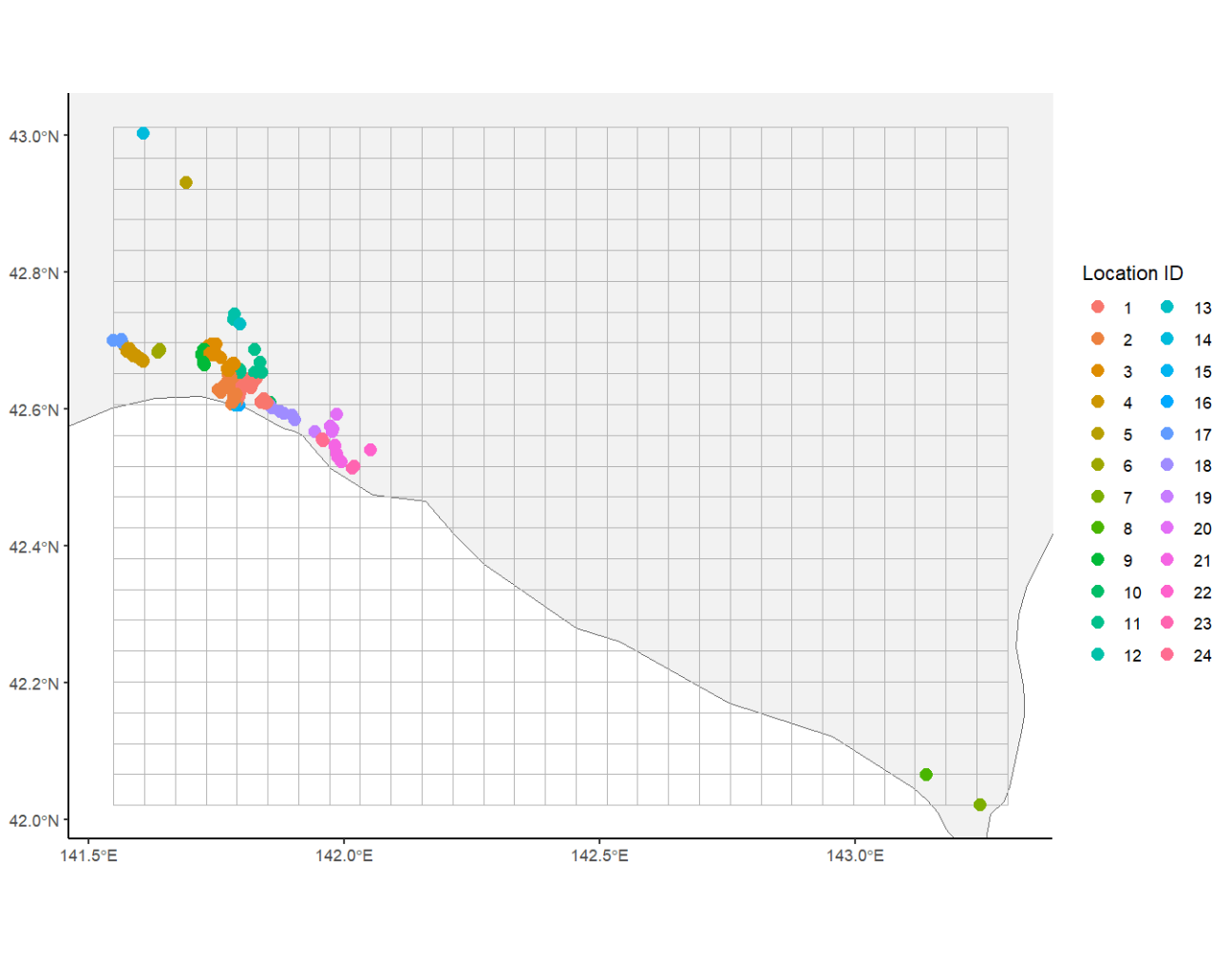

**Figure. S4.** Location IDs before excluding trials with FID > 150 m. Each grid cell represents 5 km × 5 km.

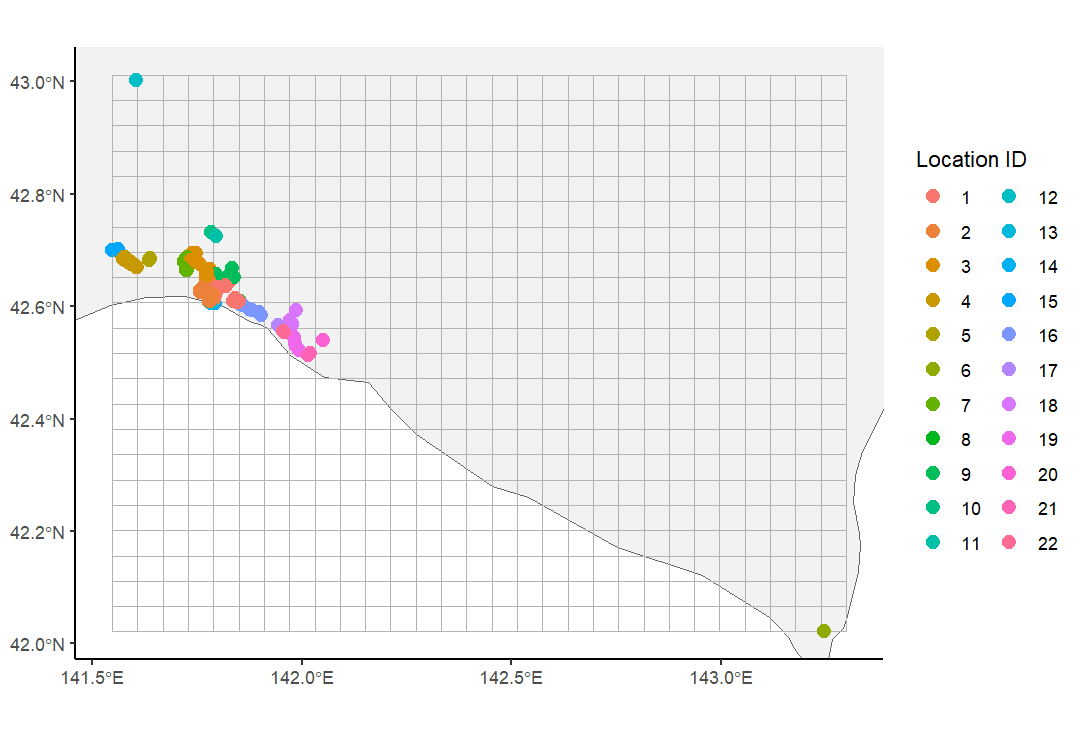

**Figure S5.** Location IDs after excluding trials with FID > 150 m. Each grid cell represents 5 km × 5 km.

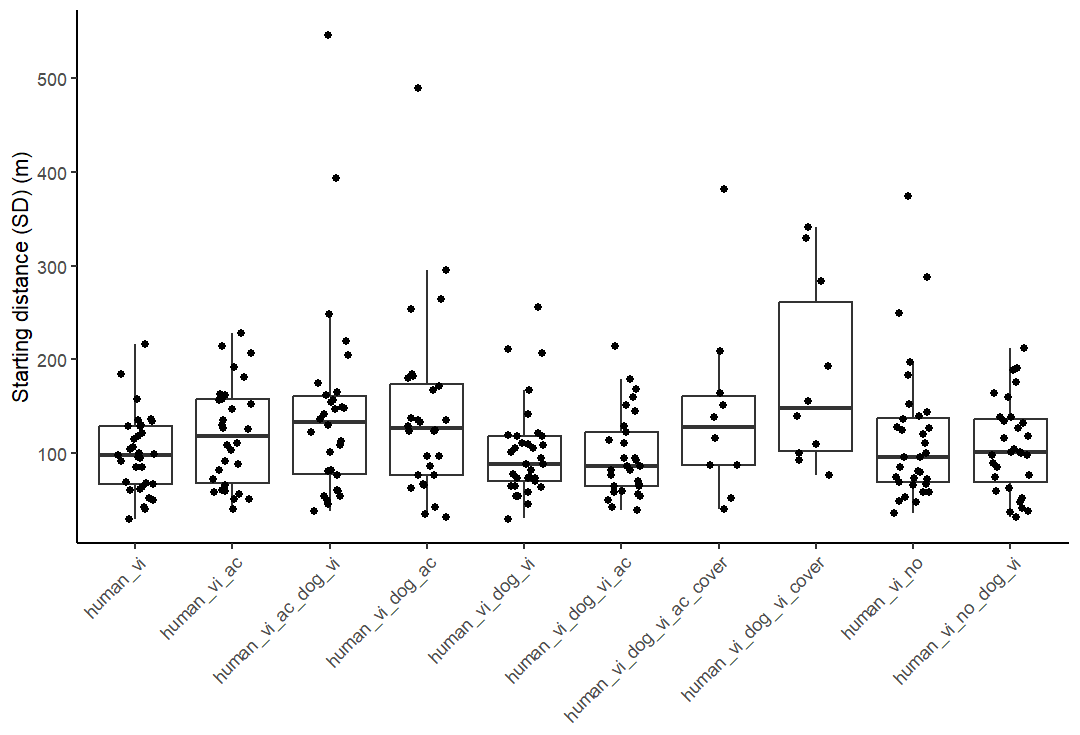

**Figure S6.** Starting distance (SD) in each trial by cue combination. The x-axis shows cue combinations (human_vi, human visual cue; human_ac, human acoustic cue; dog_vi, dog visual cue; dog_ac, dog acoustic cue; _no, white noise; dog_cover, covered dog decoy), and the y-axis shows SD.

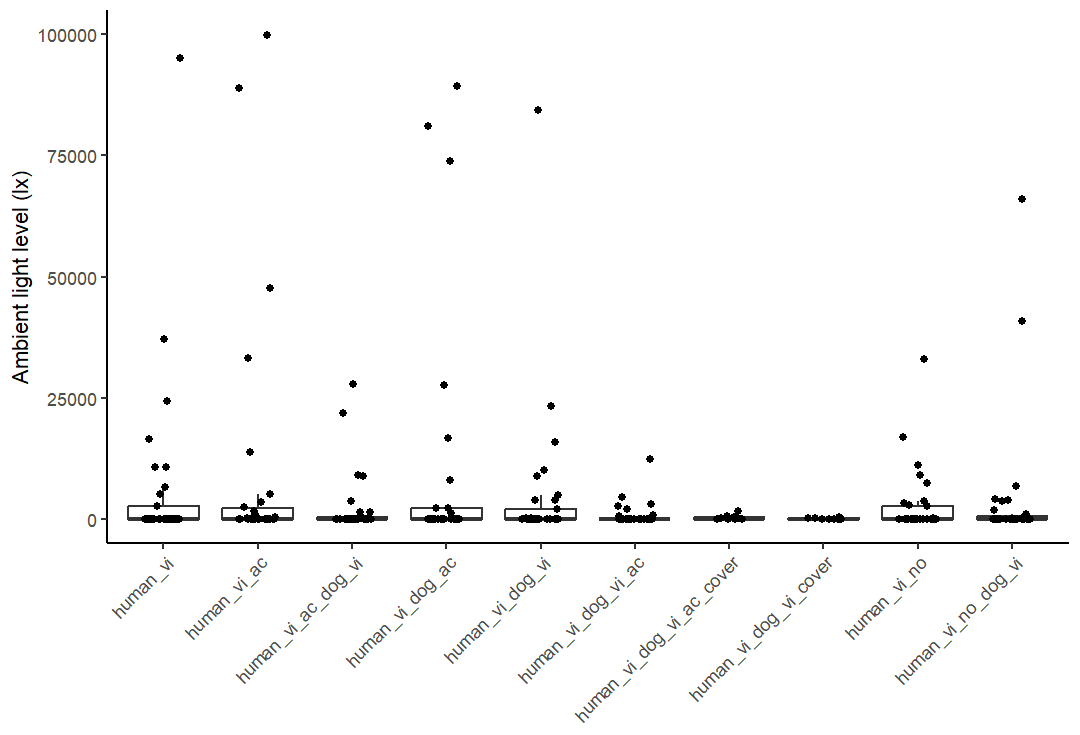

**Figure S7.** Ambient light level measured after each trial. The x-axis shows cue combination, and the y-axis shows ambient light level (lx).

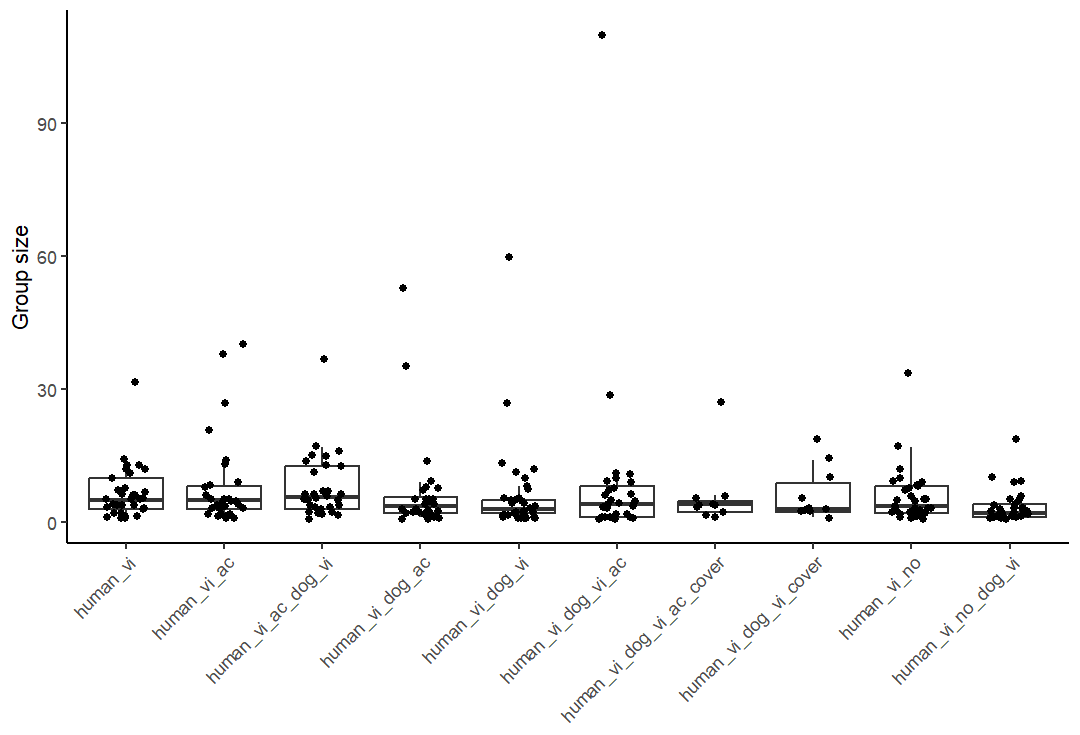

**Figure S8.** Deer group size in each trial. The x-axis shows cue combination, and the y-axis shows group size (number of individuals).

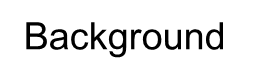

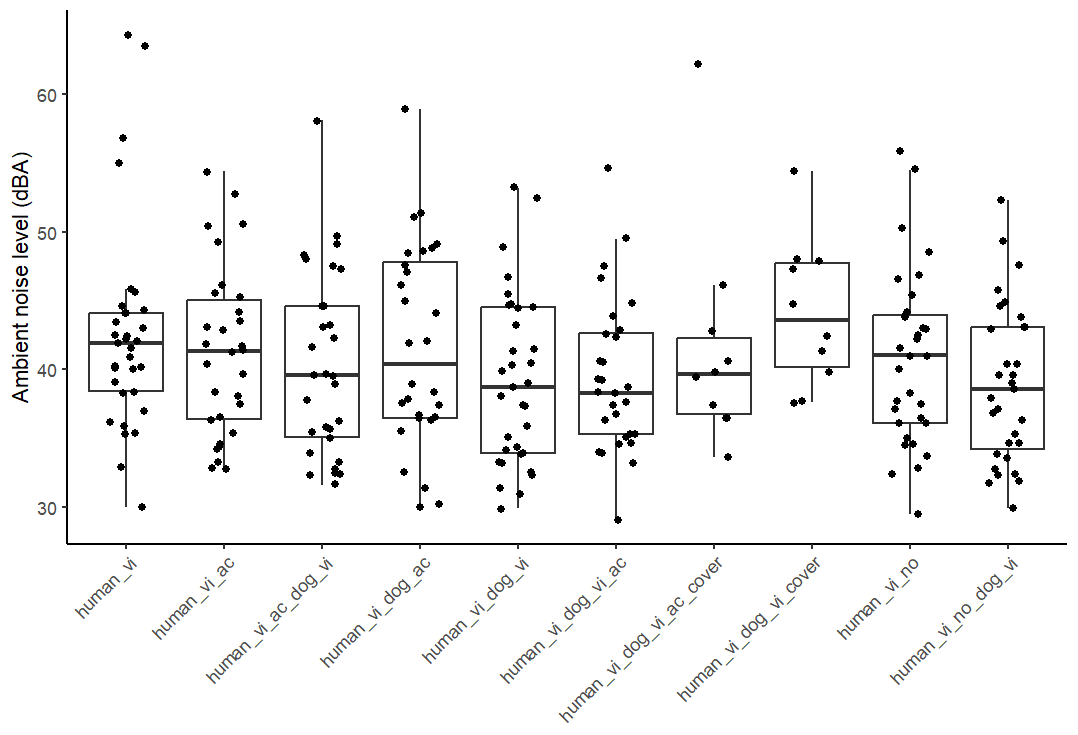

**Figure S9.** Background noise level measured after each trial. The x-axis shows cue combination, and the y-axis shows background noise level (LAeq,1min; dBA).

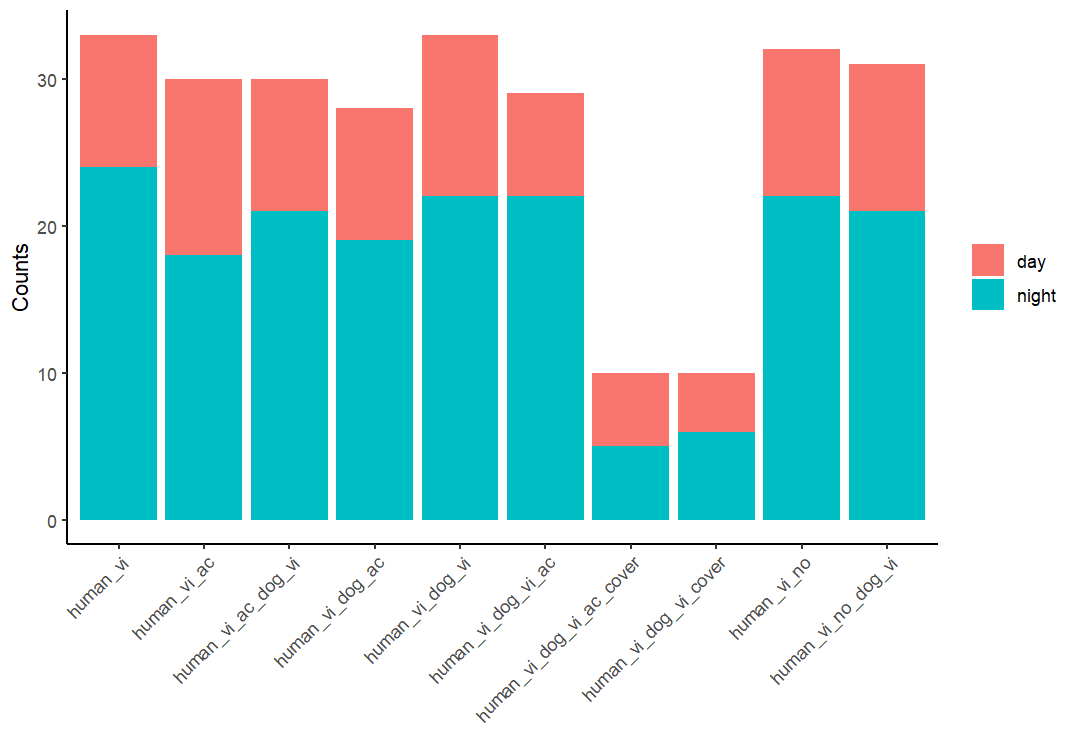

**Figure S10.** Time of day for each trial. The x-axis shows cue combination, and the y-axis shows time of day (daytime vs night-time).

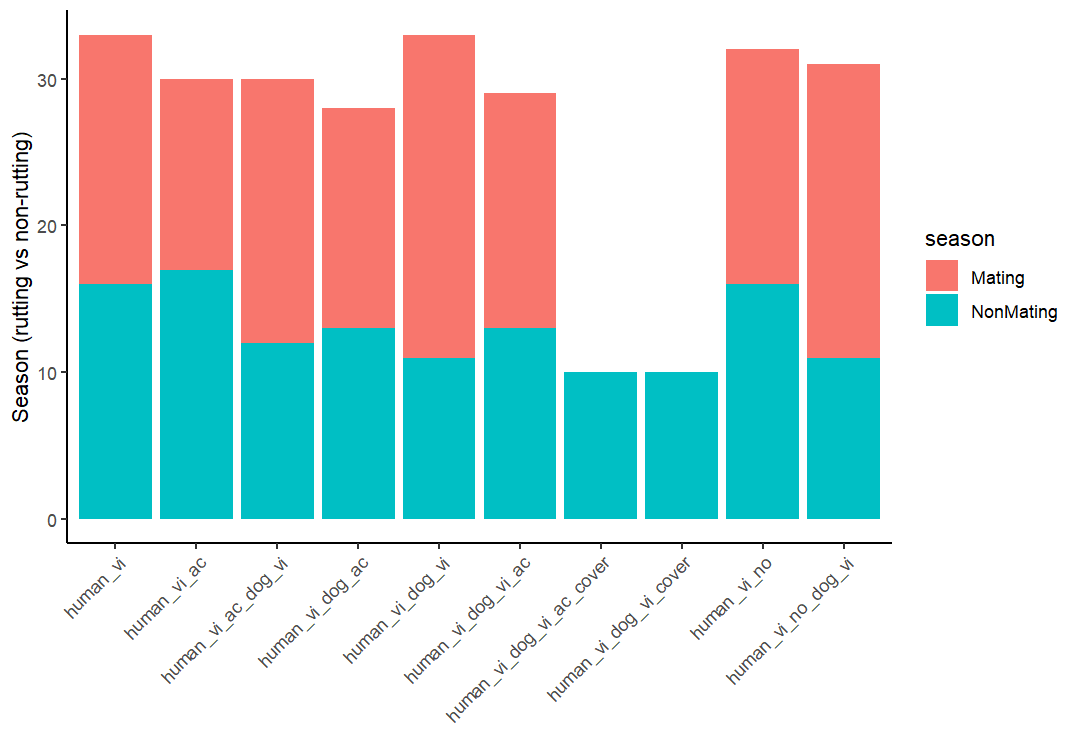

**Figure S11.** Season for each trial. The x-axis shows cue combination, and the y-axis shows season (rutting vs non-rutting).

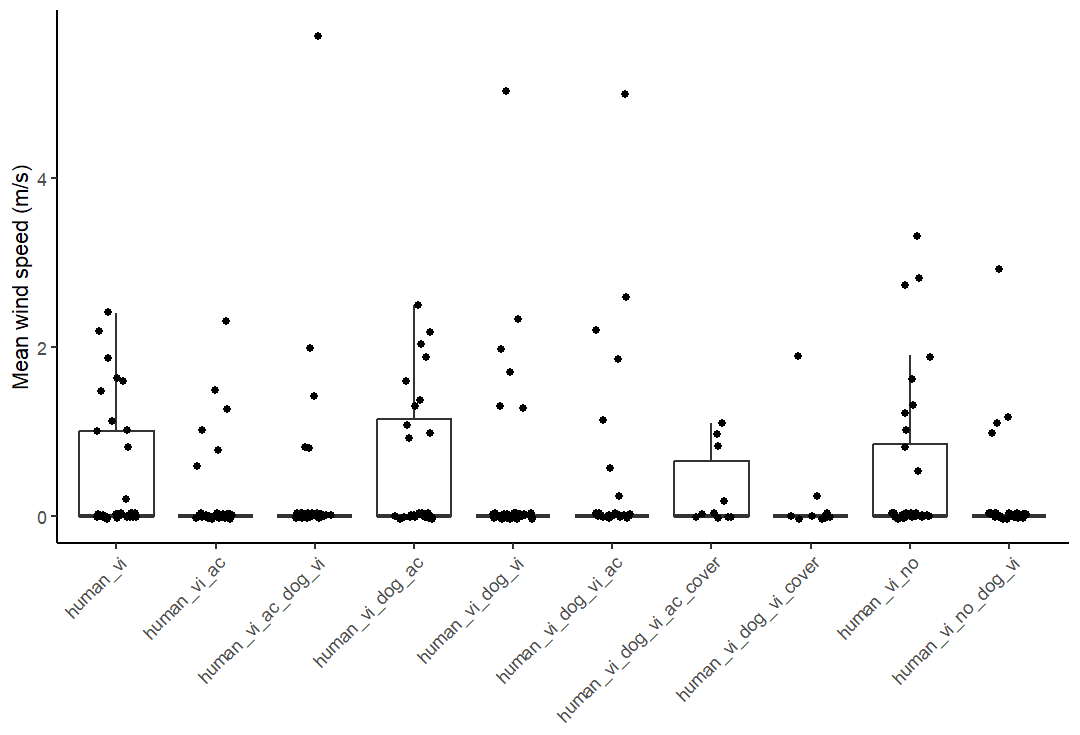

**Figure S12.** Mean wind speed measured after each trial. The x-axis shows cue combination, and the y-axis shows mean wind speed (m/s).

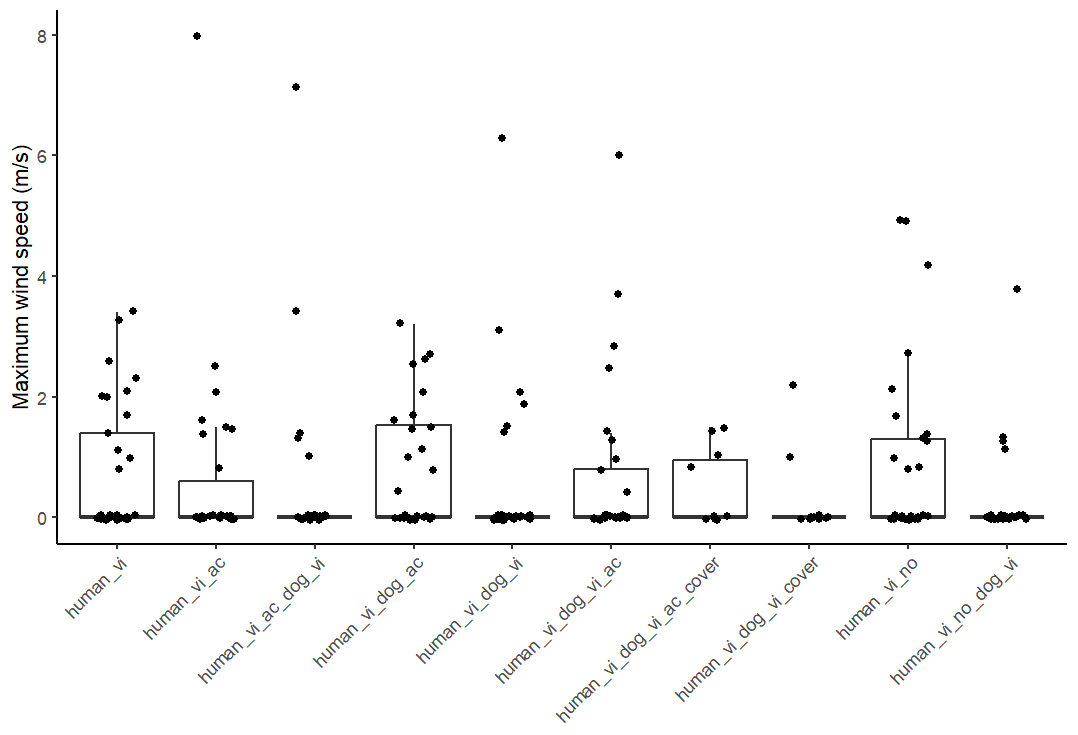

**Figure S13.** Maximum wind speed measured after each trial. The x-axis shows cue combination, and the y-axis shows maximum wind speed (m/s).

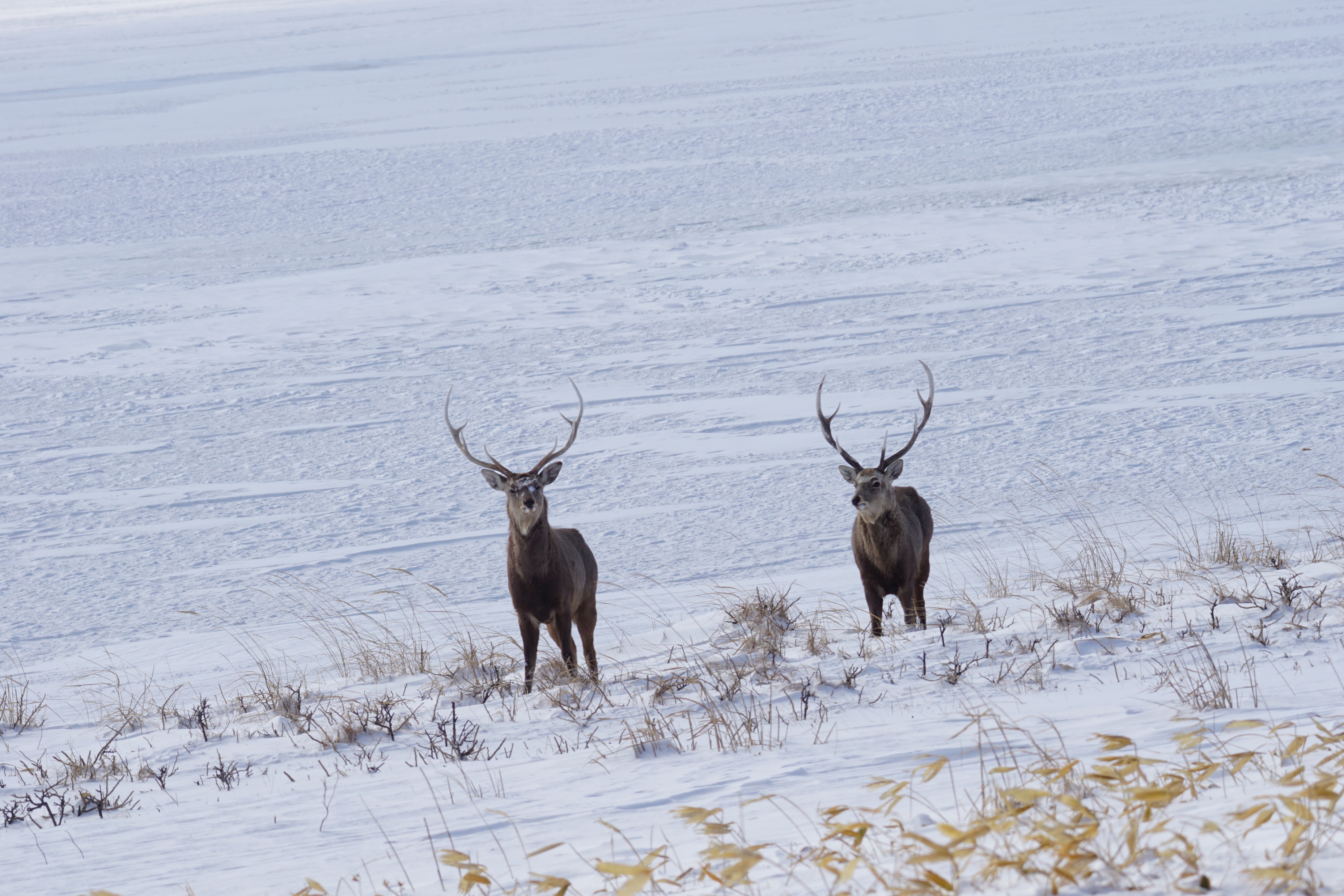

**Figure S14.** A photo of male *Cervus nippon yezoensis*. Photo credit; D. Waga.

**Appendix S1. Determining the residual error structure of the LMMs**

We could not determine from Figs S2–S3 whether the residual error structure of the LMMs should follow a Gaussian distribution or a log-normal distribution. We therefore compared model fit by calculating R² for two candidate specifications: (i) LMMs fitted with Gaussian errors on the original scale and (ii) LMMs fitted after log-transforming the response variables (AD and FID), which corresponds to a log-normal error structure on the original scale. We report results from the specification with the higher R² in the main text (Table 2). Results from the alternative (log-normal) specification are presented in Table S1.

**Appendix S2. Effect of the proportion of voiced segments in human-voice playbacks**

We fitted linear models to test whether the proportion of voiced segments in the human-voice playback files influenced AD or FID. In these models, we included relevant environmental variables as fixed effects to account for potential confounding. The results are presented in Table S2.

**Table S1.** Results of linear mixed-effects models (LMMs) evaluating the effects of cues and environmental factors on alert distance (AD) and flight initiation distance (FID) under a log-normal error structure (log-transformed responses).

| Response variable | Explanatory variable | Estimate ± SE (Lower 95% CI, Upper 95% CI) | p value | Conditional R² | Marginal R² |
| --- | --- | --- | --- | --- | --- |
| AD | Fixed factors |  |  | 0.600 | 0.593 |
|  | Intercept | 3.59 ± 0.0647 (3.47, 3.71) | <0.01 |  |  |
|  | **Human voice (presence)** | **0.155 ± 0.0770 (-1.80, 25.5)** | **0.0447** |  |  |
|  | Dog decoy (presence) | 0.0495 ± 0.0578 (-0.0669, 0.155) | 0.392 |  |  |
|  | **Dog barking (presence)** | **0.155 ± 0.0703 (0.0161, 0.287)** | **0.0282** |  |  |
|  | Dog decoy + Human voice (presence) | -0.00717 ± 0.103 (-0.203, 0.194) | 0.945 |  |  |
|  | Dog decoy + Dog barking (presence) | 0.0188 ± 0.100 (-0.169, 0.216) | 0.851 |  |  |
|  | Covered dog decoy (presence) | -0.0655 ± 0.0887 (-0.242, 0.0967) | 0.461 |  |  |
|  | **White noise (presence)** | **0.173 ± 0.0573 (0.0594, 0.0280)** | **<0.01** |  |  |
|  | Ambient light level | 0.00551 ± 0.00588 (-00.00623, 0.0155) | 0.350 |  |  |
|  | **SD** | **0.00514 ± 0.000312 (0.00467, 0.00583)** | **<0.01** |  |  |
|  | **Group size** | **0.00411 ± 0.00207 (0.000242, 0.00818)** | **0.0475** |  |  |
|  | Mean wind speed | -0.0321 ± 0.0237 (-0.0749, 0.0153) | 0.177 |  |  |
|  | Season (non-rutting) | 0.0851 ± 0.0447 (-0.00249, 0.169) | 0.0583 |  |  |
|  | Random factors |  |  |  |  |
|  | Location ID (N = 22) | 0.00199 ± 0.0446 |  |  |  |
| FID | Fixed factors |  |  | 0.462 | 0.444 |
|  | Intercept | 3.41 ± 0.0735 (3.27, 3.55) | <0.01 |  |  |
|  | Human voice (presence) | 0.165 ± 0.0860 (-0.00177, 0.329) | 0.0557 |  |  |
|  | Dog decoy (presence) | 8.05 ± 3.94 (-0.0273, 0.222) | 0.127 |  |  |
|  | Dog barking (presence) | 0.0110 ± 0.0783 (-0.140, 0.161) | 0.889 |  |  |
|  | Dog decoy + Human voice (presence) | -0.0212 ± 0.115 (-0.242, 0.201) | 0.854 |  |  |
|  | Dog decoy + Dog barking (presence) | 0.119 ± 0.112 (-0.0950, 0.334) | 0.289 |  |  |
|  | Covered dog decoy (presence) | -0.0900 ± 0.0992 (-0.282, 0.0990) | 0.365 |  |  |
|  | White noise (presence) | 0.00533 ± 0.0639 (-0.120, 0.128) | 0.934 |  |  |
|  | Ambient light level | 0.00232 ± 0.00665 (-0.0114, 0.0152) | 0.727 |  |  |
|  | **SD** | **0.00410 ± 0.000351 (0.00343, 0.00480)** | **<0.01** |  |  |
|  | Group size | 0.00439 ± 0.00231 (-0.0000100, 0.00885) | 0.0578 |  |  |
|  | **Mean wind speed** | **-0.0595 ± 0.0266 (-0.111, -0.00883)** | **0.0259** |  |  |
|  | **Season (non-mating)** | **0.109 ± 0.0500 (-0.0115, 0.204)** | **0.0304** |  |  |
|  | Random factors |  |  |  |  |
|  | Location ID (N = 22) | 0.00449 ± 0.0670 |  |  |  |

We included location ID as a random effect to account for potential spatial autocorrelation in the dataset. Results are highlighted in bold when the 95% confidence interval does not include zero. N = 266.

**Table S2.** Results of linear models evaluating the effects of the proportion of voiced segments in human-voice playbacks on alert distance (AD) and flight initiation distance (FID), while controlling for environmental variables (fixed effects).

| Response variable | Explanatory variable | Estimate ± SE | p value |
| --- | --- | --- | --- |
| AD | Fixed factors |  |  |
|  | Intercept | 76.0 ± 32.4 | 0.0226 |
|  | Ambient light level | 0.771 ± 1.19 | 0.5209 |
|  | SD | 0.386 ± 0.0544 | <0.01 |
|  | Group size | 0.829 ± 0.526 | 0.121 |
|  | Mean wind speed | -2.23 ± 5.24 | 0.672 |
|  | Season (non-rutting) | 15.1 ± 8.93 | 0.0958 |
|  | Proportion of voiced segments | -12.1 ± 9.27 | 0.1973 |
| FID | Fixed factors |  |  |
|  | Intercept | 48.0 ± 23.4 | 0.0447 |
|  | Ambient light level | -0.310 ± 0.861 | 0.721 |
|  | SD | 0.278 ± 0.0392 | <0.01 |
|  | Group size | 0.798 ± 0.379 | 0.0400 |
|  | Mean wind speed | -3.57 ± 3.78 | 0.350 |
|  | Season (non-mating) | 15.1 ± 6.44 | 0.0230 |
|  | Proportion of voiced segments | -6.00 ± 6.69 | 0.374 |

We used AD and FID observations from trials in which we presented human voice. N = 60.

**Table S3.** Sensitivity analysis of linear models (LMs) under a normal error structure (log-transformed responses) for alert distance (AD) and flight initiation distance (FID) under unimodal and multimodal cue conditions. Results are presented in the following order: human unimodal (visual-only) AD, human multimodal (visual + acoustic) AD, dog unimodal (visual-only) AD, dog multimodal (visual + acoustic) AD; followed by the corresponding models for FID in the same order.

| Response variable | Explanatory variable | Estimate ± SE | p value |
| --- | --- | --- | --- |
| AD | Fixed factors |  |  |
|  | Intercept | 9.44 ± 9.78 | 0.343 |
|  | Ambient light level | -0.190 ± 1.07 | 0.860 |
|  | **SD** | **0.659 ± 0.0982** | **<0.01** |
|  | **Group size** | **1.04 ± 0.565** | **0.0769** |
|  | **Mean wind speed** | **-10.4 ± 4.68** | **0.0353** |
|  | Season (non-rutting) | -7.06 ± 6.68 | 0.300 |
| AD | Fixed factors |  |  |
|  | Intercept | 8.97 ± 7.53 | 0.238 |
|  | Human voice (presence) | 5.92 ± 5.77 | 0.309 |
|  | **Ambient light level** | **1.40 ± 0.774** | **0.0751** |
|  | **SD** | 0.589 ± 0.0615 | **<0.01** |
|  | **Group size** | **1.19 ± 0.367** | **<0.01** |
|  | **Mean wind speed** | **-9.96 ± 4.60** | **0.0347** |
|  | Season (non-mating) | -3.88 ± 5.77 | 0.504 |
| AD | Fixed factors |  |  |
|  | **Intercept** | **34.5 ± 14.5** | **0.0243** |
|  | Ambient light level | -0.260 ± 1.50 | 0.863 |
|  | **SD** | **0.400 ± 0.108** | **<0.01** |
|  | Group size | -0.173 ± 0.509 | 0.737 |
|  | Mean wind speed | -2.48 ± 5.10 | 0.631 |
|  | Season (non-mating) | 8.33 ± 12.1 | 0.496 |
| AD | Fixed factors |  |  |
|  | Intercept | 14.5 ± 9.50 | 0.132 |
|  | **Dog barking (presence)** | **15.7 ± 6.21** | **0.0141** |
|  | Ambient light level | -0.0728 ± 0.927 | 0.938 |
|  | **SD** | **0.578 ± 0.0704** | **<0.01** |
|  | Group size | 0.107 ± 0.205 | 0.605 |
|  | Mean wind speed | -1.34 ± 2.93 | 0.650 |
|  | Season (non-mating) | 5.88 ± 6.91 | 0.399 |
| FID | Fixed factors |  |  |
|  | Intercept | 4.42 ± 9.24 | 0.636 |
|  | Ambient light level | -0.991 ± 1.01 | 0.334 |
|  | **SD** | **0.522 ± 0.0928** | **<0.01** |
|  | Group size | 0.831 ± 0.534 | 0.132 |
|  | Mean wind speed | -6.12 ± 4.43 | 0.178 |
|  | Season (non-rutting) | 2.63 ± 6.32 | 0.680 |
| FID | Fixed factors |  |  |
|  | **Intercept** | **10.3 ± 5.91** | **0.0867** |
|  | Human voice (presence) | 1.18 ± 4.53 | 0.796 |
|  | Ambient light level | 0.0331 ± 0.608 | 0.957 |
|  | **SD** | **0.399 ± 0.0483** | **<0.01** |
|  | **Group size** | **1.19 ± 0.288** | **<0.01** |
|  | Mean wind speed | -5.81 ± 3.61 | 0.114 |
|  | Season (non-mating) | 4.22 ± 4.53 | 0.356 |
| FID | Fixed factors |  |  |
|  | **Intercept** | **24.4 ± 10.3** | **0.0253** |
|  | Ambient light level | 0.585 ± 1.07 | 0.588 |
|  | **SD** | **0.330 ± 0.0769** | **<0.01** |
|  | Group size | -2.68 ± 0.363 | 0.467 |
|  | **Mean wind speed** | **-6.33 ± 3.63** | **0.0927** |
|  | Season (non-mating) | 2.33 ± 8.60 | 0.788 |
| FID | Fixed factors |  |  |
|  | Intercept | 7.35 ± 8.24 | 0.377 |
|  | **Dog barking (presence)** | **12.6 ± 5.39** | **0.0230** |
|  | Ambient light level | 0.394 ± 0.804 | 0.626 |
|  | **SD** | **0.438 ± 0.0611** | **<0.01** |
|  | Group size | 0.140 ± 0.178 | 0.436 |
|  | Mean wind speed | -2.33 ± 2.55 | 0.364 |
|  | Season (non-mating) | 10.1 ± 6.00 | 0.0986 |

Results are highlighted in bold when p < 0.10.

**Movie S1.** Timeline of a representative trial. At 0:00, the surveyor detected a non-alert deer group and began to approach. At 0:08, the closest individual became vigilant. At 0:39, all deer started fleeing (galloping and bounding).

**Movie S2.** Timeline of an excluded trial. At 0:00, the surveyor detected a deer group, but some individuals were already alert; we excluded trials like this from the analyses. At 0:12, the closest individual began trotting, which does not meet our definition of fleeing. At 0:28, all deer started fleeing (galloping and bounding).
